# High concordance between hippocampal transcriptome of the intraamygdala kainic acid model and human temporal lobe epilepsy

**DOI:** 10.1101/2020.05.12.092338

**Authors:** Giorgia Conte, Alberto Parras, Mariana Alves, Ivana Ollà, Laura de Diego-Garcia, Edward Beamer, Razi Alalqam, Alejandro Ocampo, Raúl Mendez, David C. Henshall, José J. Lucas, Tobias Engel

## Abstract

**Objective:** Pharmacoresistance and the lack of disease-modifying actions of current anti-seizure drugs persist as major challenges in the treatment of epilepsy. Experimental models of chemoconvulsant-induced status epilepticus remain the models of choice to discover potential anti-epileptogenic drugs but doubts remain as to the extent to which they model human pathophysiology. The aim of the present study was to compare the molecular landscape of the intraamygdala kainic acid model of status epilepticus in mice with findings in resected brain tissue from patients with drug-resistant temporal lobe epilepsy (TLE).

**Methods:** Status epilepticus was induced via intraamygdala microinjection of kainic acid in C57BL/6 mice and gene expression analysed via microarrays in hippocampal tissue at acute and chronic time-points. Results were compared to reference datasets in the intraperitoneal pilocarpine and intrahippocampal kainic acid model and to human resected brain tissue (hippocampus and cortex) from patients with drug-resistant TLE.

**Results:** Intraamygdala kainic acid injection in mice triggered extensive dysregulation of gene expression which was ∼3-fold greater shortly after status epilepticus (2729 genes) when compared to epilepsy (412). Comparison to samples of patients with TLE revealed a particular high correlation of gene dysregulation during established epilepsy. Pathway analysis found suppression of calcium signalling to be highly conserved across different models of epilepsy and patients. CREB was predicted as one of the main up-stream transcription factors regulating gene expression during acute and chronic phases and inhibition of CREB reduced seizure severity in the intraamygdala kainic acid model.

**Significance:** Our findings suggest the intraamygdala kainic acid model faithfully replicates key molecular features of human drug-resistant temporal lobe epilepsy and provides potential rationale target approaches for disease-modification through new insights into the unique and shared gene expression landscape in experimental epilepsy.

**Key point box:**

- More genes show expression changes shortly following intraamygdala kainic acid-induced status epilepticus when compared to established epilepsy.
- The intraamygdala kainic acid mouse model mimics closely the gene expression landscape in the brain of patients with temporal lobe epilepsy.
- Supressed calcium signalling in the brain as common feature across experimental models of epilepsy and patients with temporal lobe epilepsy.
- CREB is a major up-stream transcription factor during early changes following status epilepticus and once epilepsy is established.

## Introduction

A major challenge in epilepsy is the lack of adequate treatment with >30% of patients remaining resistant to currently available anti-seizure drugs (ASDs).^1, 2^ Moreover, ASDs may cause severe adverse-effects and, critically, current pharmacological treatment remains purely symptomatic and does not significantly alter the course of the disease.^3^ Thus, there is a pressing need for the identification of drug targets with a different mechanism of action. The most common drug refractory form of epilepsy in adults is temporal lobe epilepsy (TLE) involving different structures within the limbic system, including the amygdala and hippocampus.^4^ Pathological processes occurring during epileptogenesis include structural and functional changes such as ongoing neurodegeneration and reorganization of neural networks.^5^ Mounting data obtained via gene expression profiling suggests these processes are driven in part by large-scale changes in the gene expression landscape within the brain.^6-14^

Whereas animal models of acute seizures (*e.g.*, pentylenetetrazol and maximal electroshock) have been successful in the identification of ASDs, the identification of anti-epileptogenic drugs and drugs to treat refractory epilepsy most likely requires different models that phenocopy the chronic stage of the disease.^15^ The translational value of a model depends, however, on there being extensive homology with the human pathophysiology or we risk developing ineffective treatments. Currently, rodent models of status epilepticus (SE) remain the preferred method for generating drug-resistant epilepsy. The extent to which these match the molecular landscape of human refractory TLE remains uncertain. The intraamygdala kainic acid (IAKA) model of focal-onset SE in mice produces unilateral pathology and drug-resistant epilepsy after a short latent period.^16^ The model is increasingly used for studying mechanisms of epileptogenesis and the testing of novel anti-epileptogenic drugs.^17-23^ Whereas changes in the transcriptome have been analysed shortly following SE in the model,^24, 25^ global changes in gene expression during chronic epilepsy and whether alterations in gene expression reflect changes occurring in patients remains to be established.

Here, we compared the gene expression profile in the IAKA mouse model with reference data from patients with drug-resistant TLE and two other models of SE as comparator. Our results reveal excellent molecular concordance between the IAKA model and human TLE and identify potential targets for disease-modifying treatments.

## Methods

### Animal model of status epilepticus

Animal experiments were performed in accordance with the principles of the European Communities Council Directive (2010/63/EU) and approved by the Research Ethics Committee of the Royal College of Surgeons in Ireland (RCSI) (REC 1322, 842) and the Irish Health Products Regulatory Authority (AE19127/P038). Experiments were carried out in 8-12-week-old C57Bl/6 male mice bred at RCSI.^26^ SE was induced in fully awake mice via a microinjection of kainic acid (KA) (0.3 µg in 0.2 µl phosphate-buffered saline (PBS)) (Sigma-Aldrich, Dublin, Ireland) into the basolateral amygdala. Vehicle-injected control animals received 0.2 µl PBS. The anticonvulsive lorazepam (6mg/kg) (Wyetch, Taplow, UK) was delivered i.p. 40 min post-IAKA or vehicle to curtail seizures and reduce morbidity and mortality. Electroencephalogram (EEG) was recorded from cortical implanted electrodes (Xltek recording system; Optima Medical Ltd, Guildford, UK) starting 10 min prior IAKA administration.

### EEG quantification and behavioral assessment of seizure severity during status epilepticus

Seizures were analysed via Labchart7 (ADInstruments Ltd, Oxford, United Kingdom).^27^ EEG total power (μV^2^) was analysed by integrating frequency bands from 0-100 Hz. Power spectral density heat maps were generated using Labchart7 (frequency (0-40 Hz), amplitude (0-50 mV)). Clinical behaviour were scored every 5 min for 40 min after IAKA according to a modified Racine Scale.^28^ Score 1, immobility and freezing; Score 2, forelimb and/or tail extension, rigid posture; Score 3, repetitive movements, head bobbing; Score 4, rearing and falling; Score 5, continuous rearing and falling; Score 6, severe tonic– clonic seizures. The highest score attained during each 5 min period was recorded by an observer blinded to treatment.

### Drug treatment

To determine the effects of CREB1 on SE, additional mice received an intracerebroventricular (i.c.v.) infusion of 2 nmol of the CREB1 inhibitor 666-15^29, 30^ in 2 μl volume (PBS in 0.2% Dimethyl sulfoxide) 10 min before IAKA and 60 min post-lorazepam treatment to reach a final concentration of 1 mM in the ventricle (ventricle volume was calculated as 35 µl).

### RNA isolation and microarrays analysis

Mice were sacrificed 8 h or 14 days post-SE. Ipsilateral hippocampi were quickly dissected and pooled (n = 3 per pooled sample). Total tissue RNA was extracted using the Maxwell® 16 LEV simplyRNA Tissue Kit (Promega, AS1280). RNA quantification was performed with Qubit RNA Hs Assay kit (Thermo-Fisher Scientific, Q32852) and RNA integrity QC with Agilent Bioanalyzer 2100, using RNA Nano Assay (Agilent Technologies 5067-1511) and RNA Pico Assay (Agilent Technologies 5067-1513). cDNA library preparation and amplification were performed with WTA2 kit (Sigma-Aldrich) using 2-5 ng of total RNA as template. cDNA was amplified for 22 cycles and purified using PureLink Quick PCR Purification Kit (Invitrogen, K310001). 8 µg of the cDNA from each sample were fragmented and labelled with GeneChip Mapping 250K Nsp assay kit (Affymetrix, 900753). Hybridization was performed as described previously^31^ during 16 h at 45°C. Washing and staining steps were performed in the GeneAtlas Fluidics Station (Affymetrix, 00-0079), following the specific script for Mouse MG-430 PM Arrays. Arrays were scanned with GeneAtlas Scanner, and CEL files were done with GeneAtlas software (Affymetrix). Processing of microarray samples was performed using R and Bioconductor.^32^ Raw CEL files were normalized using RMA background correction and summarization. Probeset annotation was performed using Affymetrix version na35. For each gene, a linear model was used to find significant differences between IAKA- or vehicle (PBS)-treated mice. Analysis of differential expression was performed using a linear model implemented in the R package “limma”^33^. *P*-values were adjusted with the Benjamini and Hochberg correction. We considered one gene to be down-regulated with an adjusted *P*-value ≤ 0.01 and FC < -1.2 and up-regulated with adjusted *P*-value ≤ 0.01 and FC > 1.2 in at least one probe. If the same transcript showed opposite results for different probes, the transcript was considered as not changed.

### RNA extraction and quantitative PCR (qPCR)

RNA extraction was performed using whole hippocampi.^34^ RNA concentration was measured via a nanodrop Spectrophotometer (Thermo Scientific, Rockford, IL, U.S.A) and absorbance determined at a ratio of 260/280. Samples with an absorbance ratio between 1.8-2.2 were considered acceptable. 500 µg of total mRNA was used to produce cDNA by reverse transcription using SuperScript III reverse transcriptase enzyme (Invitrogen, CA, U.S.A). qPCR was performed using LightCycler 1.5 (Roche Diagnostics, GmbH, Mannheim, Germany). Each reaction tube contained 2 μl cDNA, 10 μl SyBR green Quantitect Regent (Quiagen Ltd, Hilden, Germany), 1.25 μM primer pair (Sigma, Dublin, Ireland) and RNAse free water (Invitrogen CA, U.S.A) to a final volume of 20 μl. Data were analysed and normalized to the expression of *β-actin*. Primers used (Sigma, Dublin, Ireland) are listed in **Table S4**.

### Comparison of gene expression changes between models

Overlap of differentially expressed genes between different models was analyzed using the hypergeometric distribution test. Gene profiling data of the pilocarpine mouse model (12 h and 6 weeks post-SE) was obtained from^10^ and for the intrahippocampal KA (IHKA) model (6 h and 2 weeks post-SE) from.^35^ Transcriptome changes in TLE patients was previously reported in^14^ for cortical tissue samples from 86 mesial temporal lobe epilepsy (mTLE) patients vs. 75 neurologically healthy controls, and^12^, for sclerotic hippocampus vs. non-spiking neocortex from 10 TLE patients. Genes were considered overlapping when the representation factor (RF) was > 2 and *P* < 0.05, and dissimilar when RF was < 0.5 and P < 0.05. Complete set of genes used in our study is shown in **Table S2**.

### Gene Ontology analysis

Pathways analysis of up- and down-regulated genes (adj. *P*-value ≤ 0.01 and FC > ± 1.2) in the IAKA model (8 h and 14 days post-SE) and of overlapping genes between IAKA model and TLE patients^12^ were analyzed using DAVID Bioinformatics Resources 6.7, KEGG pathway annotation.^36^ This included also the analysis of Calcium signalling pathways, down-regulated transcripts in TLE patients and different epilepsy mouse models.

### IPA analysis

Analysis of upstream regulators was performed applying QIAGEN’s Ingenuity® Pathway Analysis (IPA®) (QIAGEN Inc., www.qiagen.com/ingenuity)^37^ to genes found differentially expressed (adj. P-value < 0.01 and FC > 1.5) in the IAKA mouse model at 8 h and 14 days post-SE.

### Data analysis

Statistical analysis was performed using SPSS 25.0 (SPSS® Statistic IBM®). Data are represented as mean ± S.E.M. (Standard Error of the Mean) with 95% confidence interval. Higher or lower points (outliers) are plotted individually or are not plotted. The normality of the data was analyzed by Shapiro-Wilk test (n < 50) or o Kolmogorov-Smirnov (n > 50). Homogeneity of variance was analyzed by Levente test. For comparison of two independent groups, two-tailed unpaired t-Student’s test (data with normal distribution), Mann-Whitney-Wilcoxon or Kolmogorov-Smirnov tests (non-normal distribution) was performed. Enrichment tests were carried out by using one-sided Fisher’s exact test. Significance was accepted at *P* < 0.05.

### Data availability

Data supporting the findings of this study are available from the corresponding authors upon reasonable request. Records have been approved and assigned GEO accession numbers; however, until the acceptance of the manuscript, these will be kept private. GSE122228 - Identification of differentially expressed genes in the hippocampus in the intraamygdala kainic acid mouse model of SE.

## Results

### Larger transcriptome changes following status epilepticus compared to established epilepsy in intraamygdala KA-treated mice

To study global changes in transcript levels following SE and during epilepsy, mRNA was extracted from ipsilateral hippocampi of mice subjected to IAKA-induced SE^16^ at two different time-points and analysed via microarrays (total of 21549 genes). As a first time-point we chose 8 h post-SE, time-point when EEG activity usually has returned to baseline levels and first signs of seizure-induced neurodegeneration appear; however, without wide-spread cell death observed at 24 h post-SE.^38^ As a second time-point we chose 14 days post-SE, a time-point when all mice subjected to IAKA normally experience the occurrence of spontaneous seizures.^16, 19, 25^ To reduce inter-sample variability, each sample analysed was a pool of three ipsilateral hippocampi (**Fig. 1A**). Genes were considered as differentially expressed when the adjusted *P*-value was ≤ 0.01 and fold change (FC) < -1.2 for down-regulated and FC > 1.2 for up-regulated transcripts (**Fig. 1B, Table S1**). A set of transcripts were also validated via qRT-PCR, which confirmed the microarray findings for *Hspa1b, Atf4, Ctsz* and *Laptm5* (**Fig. S1A**,**B**).

**Figure 1.**
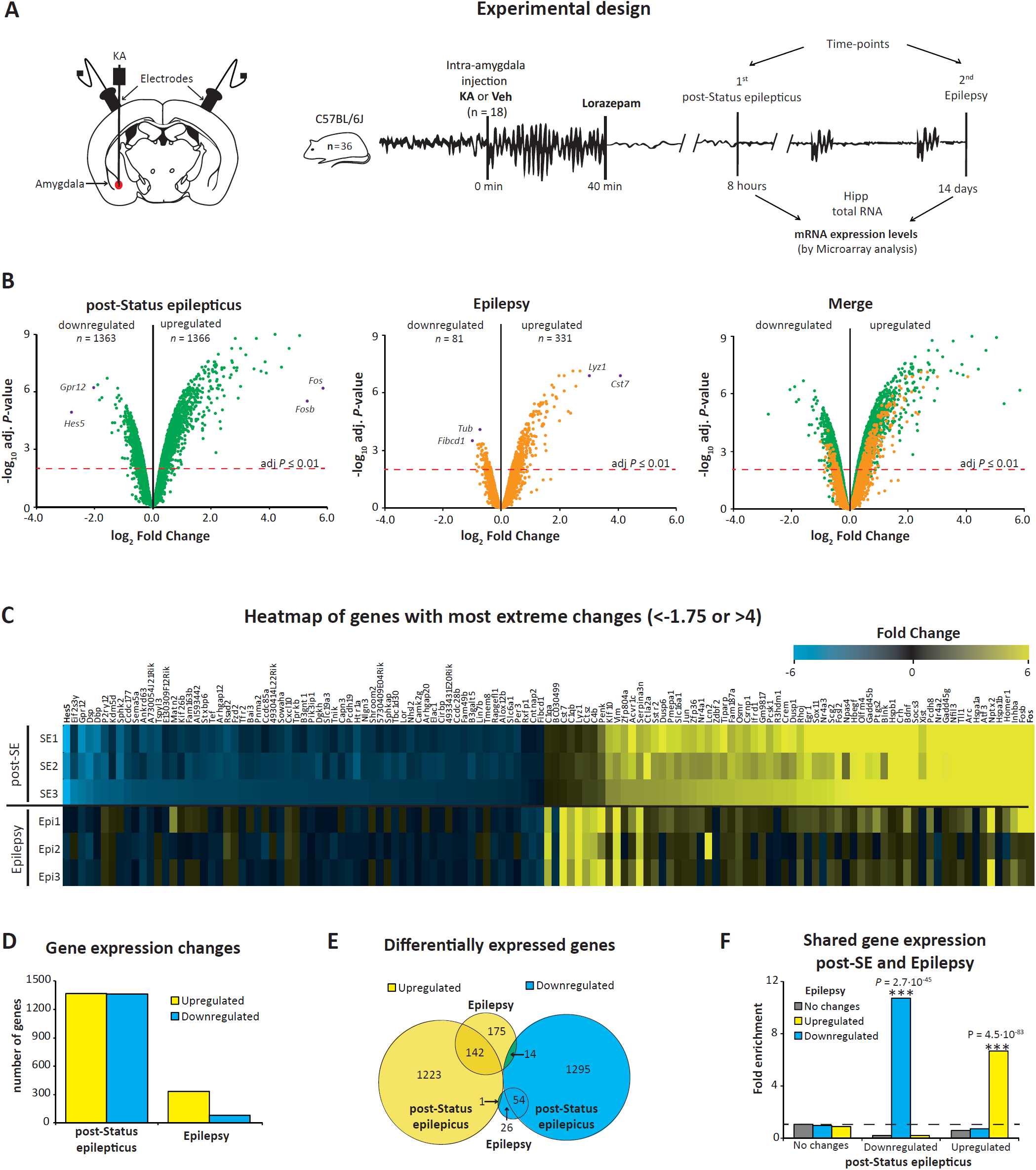
Gene expression analysis post-intraamygdala KA-induced status epilepticus and during epilepsy. (**A**) Schematic showing experimental design using the intraamygdala kainic acid (IAKA)-induced SE mouse model. Mice were euthanatized at two time-points: 8 h post-SE (acute) and 14 days (chronic epilepsy). Ipsilateral hippocampi were removed and transcript levels analyzed via microarray. (**B**) Volcano plot of genes analyzed by microarray. The X-axis represents the log_2_ ratio of gene expression levels and the Y-axis the adjusted *P*-value based on –log10. The red dashed line denotes the significant level (*P* ≤ 0.01). Purple dots represent genes with most dramatic expression changes. (**C**) Heatmap of genes showing the most dramatic expression changes (down-regulated fold change (FC) < -1.75 blue; up-regulated FC > 4 yellow), post-SE (acute) (top part) and during chronic epilepsy (bottom part). (**D**) Bar chart representing the number of dysregulated genes post-SE and in chronic epilepsy. (**E**) Venn diagram showing overlap of differentially expressed genes at the two time-points analyzed. (**F**) Comparison of differentially expressed genes post-SE and during epilepsy. The Y-axis represents the fold change enrichment and X-axis shows the different gene regulation post-SE (8 h). Colours show the different gene regulation during epilepsy (14 days). One-sided Fisher’s exact test (\*\*\**P* < 0.001).

The greatest fold changes in gene expression post-SE were found for known activity-regulated transcripts including c-*Fos* (FC = 58.1) and *Inhb* (FC = 33.2). The most down-regulated transcripts were *Hes5* (FC = -6.8) and *Gpr12* (FC = -4.0) (**Fig. 1C**). A similar number of genes were up- (1366) and down-regulated (1363) post-SE. In contrast, during epilepsy, genes were mainly up-regulated (331) with only 81 genes down-regulated (**Fig. 1D**). Furthermore, more genes showed alterations of their mRNA levels following SE (2729 genes, 12.6%) when compared to established epilepsy (412 genes, 1.9%) (**Fig. 1D,E**). Finally, a very significant overlap was found between genes up- and down-regulated in the same direction at both time-points (**Fig. 1F**).

Taken together, our results show that IAKA-induced SE leads to a unique expression profile during SE and epilepsy with changes in mRNA levels being more prominent post-SE when compared to established epilepsy.

### The intraamygdala KA mouse model mimics the hippocampal gene expression profile of TLE in humans

Chemoconvulsant-induced SE is one of the most common strategies to identify novel anti-epileptic drugs with KA and pilocarpine being the most frequently used.^15^ In order to establish whether gene expression changes are similar between models, we compared transcriptome profiles of the IAKA mouse model with transcriptome changes reported for other models were epilepsy is induced via SE including the IHKA mouse model^35^ and the i.p. pilocarpine mouse model^10^ (**Table S2**). This revealed a similar expression pattern between the IAKA and i.p. pilocarpine mouse model at both time-points, post-SE and during epilepsy, with expression changes being more similar during established epilepsy (**Fig. 2A**). Notably, the IAKA and IHKA mouse model showed an almost complete overlap in gene expression at both time-points, post-SE and epilepsy (**Fig. 2B**).

**Figure 2.**
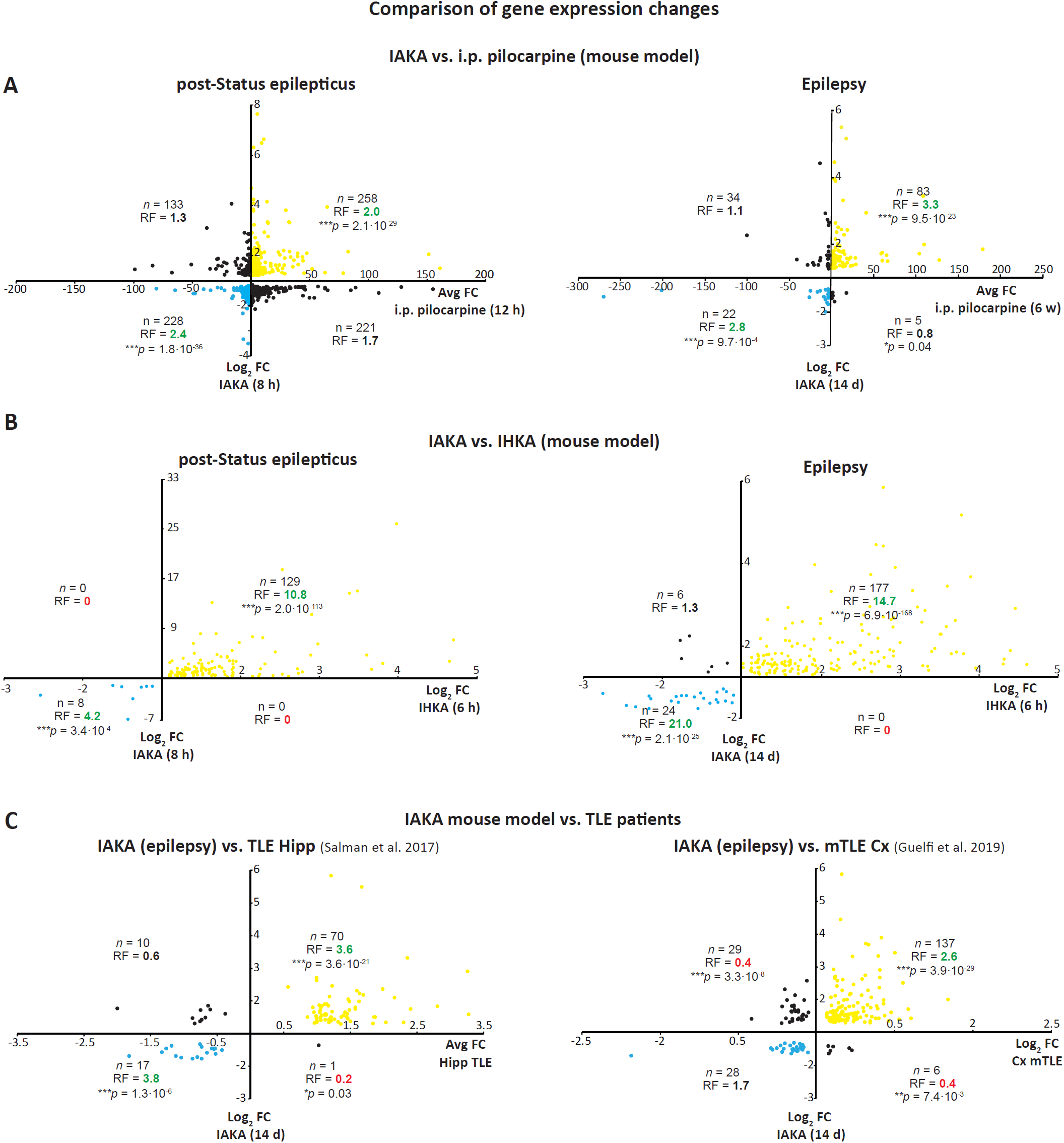
Comparison of genes undergoing expression changes in experimental models of epilepsy and TLE patients. Graphs showing comparison of dysregulated genes in the hippocampus of (**A**) mice subjected to IAKA and intraperitoneal (i.p.) pilocarpine and (**B**) mice subjected to IAKA and IHKA post-SE and during chronic epilepsy. (**C**) Comparison of dysregulated genes in hippocampus of IAKA (14 days) and hippocampi from patients with TLE (left panel) and cortex from patients with mTLE (right panel). Up-regulated genes are presented in yellow in both conditions and down-regulated genes are presented in blue. Overlap: Representation Factor (RF) > 2 and *P* < 0.05, and dissimilar RF < 0.5 and P < 0.05. Hypergeometric test.

Next, to determine the extent to which mRNA changes occurring in mouse models are translatable to TLE in humans, we compared transcriptome changes with recently published transcriptome changes in TLE patients. This included a study analysing changes in sclerotic hippocampi *vs*. neocortex of TLE patients^12^ and a study analysing gene expression changes in cortical tissue from patients with mTLE^14^ (**Table S2**). Here we found a very strong correlation between transcriptome changes identified in the IAKA mouse model during established epilepsy with transcriptome changes observed in TLE patients (hippocampus and cortex) (**Fig. 2C**). Similarly, to the IAKA model, there was also a strong correlation of dysregulated genes between the IHKA model and TLE patients (**Fig. S2A**). In contrast, the transcriptional profile differed greatly when comparing between human TLE and expression changes 8 h following IAKA-induced SE (**Fig. S2B**). Finally, we also compared transcriptome profiles between the i.p pilocarpine mouse model and TLE patients. Although there was a good overlap among down-regulated genes between pilocarpine-injected mice and TLE patients, this correlation was lost among up-regulated genes. Moreover, the magnitude of enrichments was minor when compared to both KA mouse models (**Fig. S2C**), suggesting KA mouse models reflecting closer molecular changes that occur in the brain of TLE patients.

### Suppressed calcium signalling as a common feature across experimental models of epilepsy and TLE patients

To identify pathways and functional groups of genes affected by expression changes in the IAKA mouse model, differentially expressed transcripts were analyzed by GO terms using the bioinformatic tool DAVID.^39^ This revealed that post-SE up-regulated genes are mainly associated with signalling pathways (*e.g., MAPK, PI3K-Akt*) and down-regulated genes with metabolism and synaptic transmission (**Fig. 3A** and **Table S3**). At 14 days post-SE when epilepsy was established, up-regulated genes are highly enriched in pathways linked to waste removal (*e.g., Phagosome, Lysosome*) and down-regulated genes are mainly linked to pathways controlling synaptic transmission (**Fig. 3A**), similar to down-regulated genes post-SE (**Fig. 3A, Table S3**).

**Figure 3.**
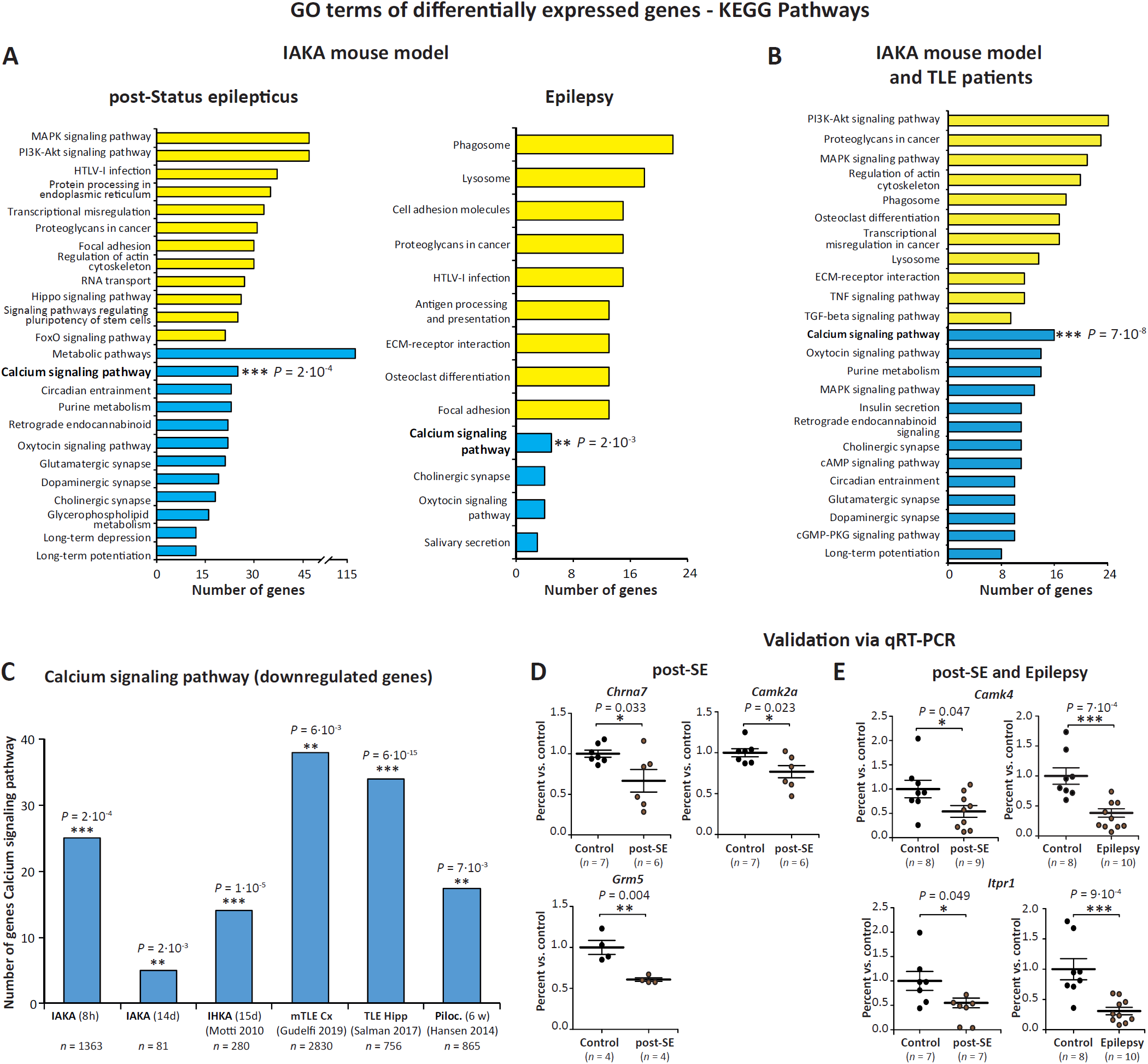
Gene Ontology (GO) analysis and validation of microarray results. Gene count histogram from GO analysis using DAVID resources of differentially expressed genes in (**A**) post-SE and in epilepsy, (**B**) TLE patients and IAKA epileptic mice common dysregulated genes. (**C**) Number of genes of calcium signalling pathway in down-regulated genes in IAKA and IHKA mouse models of epilepsy, in TLE patients (hippocampus and cortex) and in the i.p. pilocarpine model of epilepsy (one-sided Fisher’s exact test). (**D-E**) mRNA level analysis by qRT-PCR of down-regulated genes involved in calcium signalling, (**D**) following SE (control n= 4/7, post-SE n= 4/6) and (**E**) at both time-points (control n = 7 (*Ltpr1*) and 8 (*CamK4*), post-SE n = 7 (*Ltpr1*) and 9 (*CamK4*); control n = 8, epilepsy n = 10). Two-sided unpaired t-test. Data are mean ± S.E.M. * *P* < 0.05 ** *P* < 0.01 *** *P* < 0.001.

GO term analysis of dysregulated genes shared between the IAKA mouse model (established epilepsy) and TLE patients found a significant enrichment of genes linked to intracellular signalling (*e.g., PI3K-Akt, MAPK, TGF-β*) and lysosome activity among up-regulated genes and genes associated with synaptic transmission (*e.g., Calcium signalling, Long-term potentiation*) among down-regulated genes (**Fig. 3B**). Most notably, genes enriched in *Calcium signaling* were significantly affected in all three conditions (**Fig. 3A,B** and **Table S3**).

To further assess whether the enrichment of down-regulated genes involved in calcium signalling is a common response across different models, GO terms of altered transcripts were analyzed in all three animal models (KA, pilocarpine) and TLE patients.^12, 14^ This confirmed *Calcium signalling* being a common pathway suppressed in mouse models of epilepsy and TLE patients (**Fig. 3C, Table S3**). Down-regulated genes involved in *Calcium signaling* and altered post-SE and in human TLE are shown in **Table 1**. Microarray results were confirmed via single qRT-PCR analysis of selected down-regulated transcripts involved in *Calcium signalling* either unique to IAKA-induced SE (**Fig. 3D)** or common between SE and epilepsy (**Fig. 3E)**.

**Table 1.**
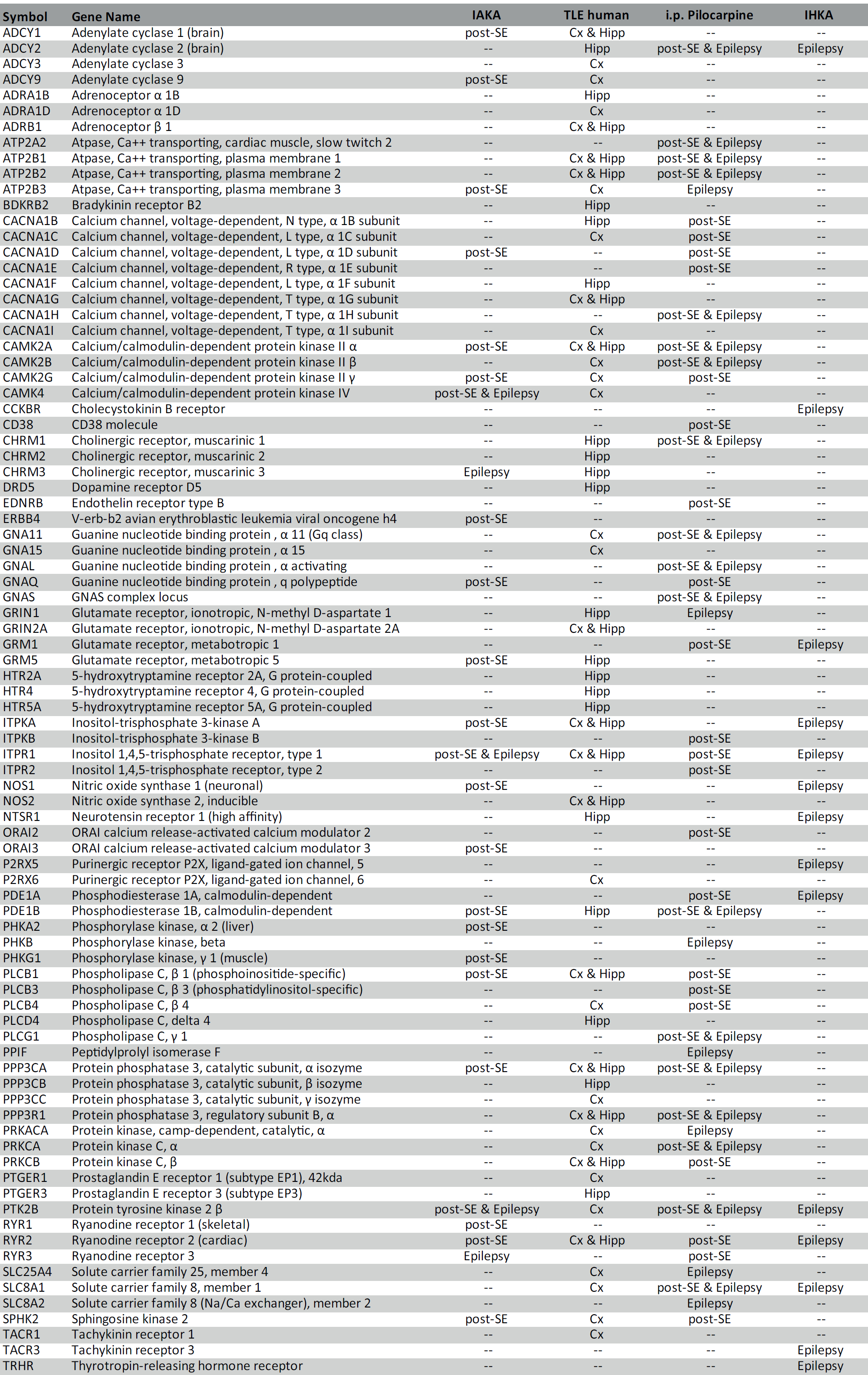
Calcium signalling (down-regulated genes) - mouse models of epilepsy and/or TLE patients

In summary, our results reveal overlap of several altered pathways between the IAKA mouse model and human TLE, particularly genes linked to calcium signaling, thereby identifying possible novel treatment targets for epilepsy.

### CREB is a major transcription factor controlling gene transcription following intraamygdala status epilepticus and during epilepsy

Analysis via the Ingenuity Pathway Analysis (IPA®)^37^ predicted several transcription factors to regulate gene expression changes post-SE and during epilepsy. Most notably, the majority of identified transcription factors are unique to SE or epilepsy (*e.g.*, TRP53 and FOS post-SE), confirming the limited overlap of differently expressed genes between conditions. Some transcription factors were, however, predicted to control gene expression in both conditions, including MYC (up-regulated genes post-SE and during epilepsy) and CREB1, linked to both up- and down-regulated genes under both conditions (**Fig. 4A**).

**Figure 4.**
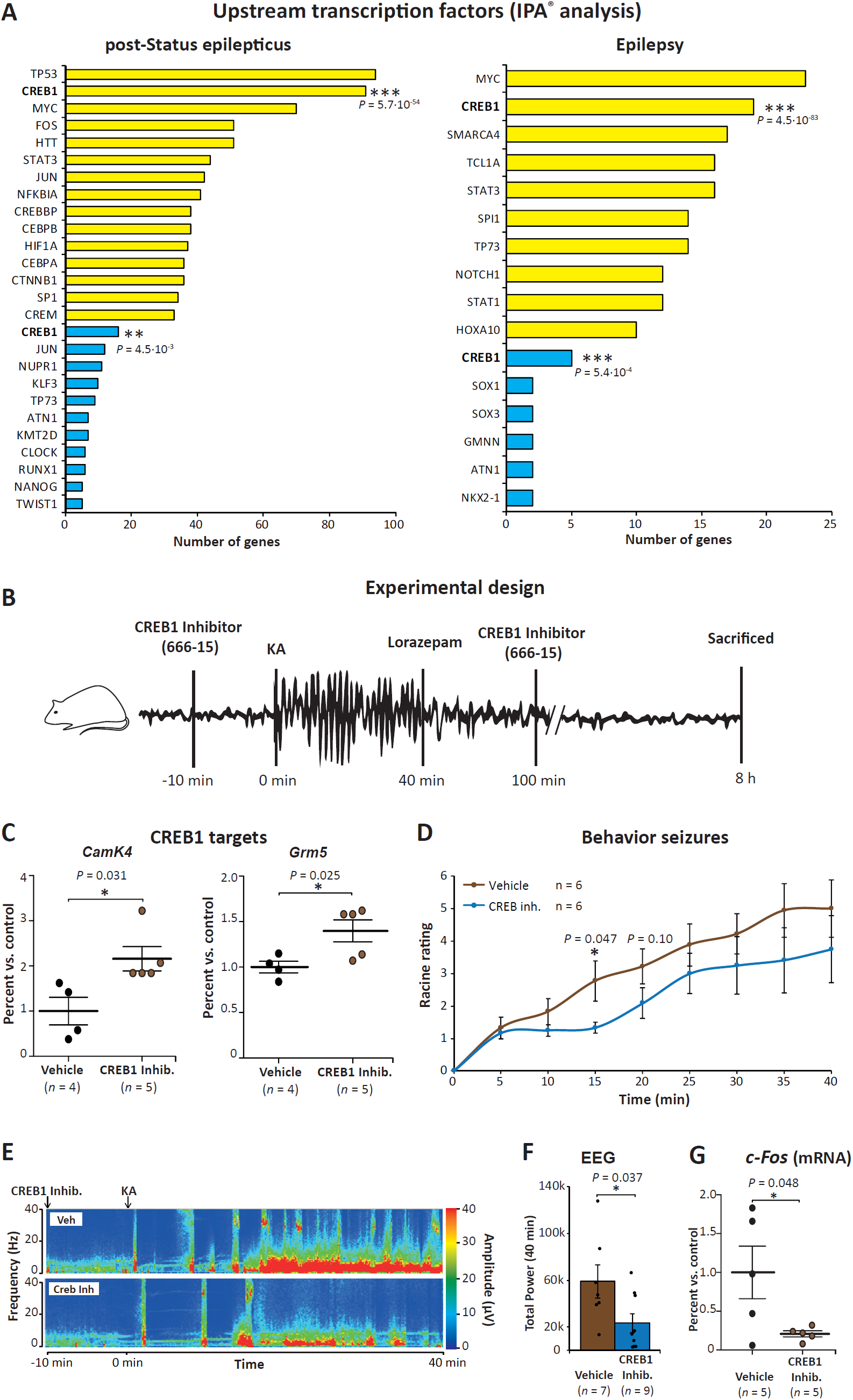
CREB1 as up-stream transcription factor during status epilepticus and epilepsy. (**A**) Prediction of transcription factors involved in gene upregulation (yellow) and downregulation (blue) post-SE and during epilepsy by using Ingenuity Pathway Analysis (IPA®). Of note, transcription factor CREB1 is one of the transcription factors with highest amount of predicted target genes among up- and down-regulated gene pool during both conditions, post-SE and during epilepsy. (**B**) Schematic of experimental design to test CREB1 inhibition on SE. Mice subjected to IAKA were treated i.c.v. with the CREB1 inhibitor 666-15 (2 nmol) 10 min before KA injection and 1 h after lorazepam treatment. (**C**) Graphs showing increased mRNA levels of the two CREB1 target genes, *CamK4* and *Grm5*. (**D**) Behavioural severity of seizures (mean Racine score) scored each 5 min and total score, in mice subjected to IAKA treated with vehicle (Veh) (n = 4) and treated with the CREB1 inhibitor 666-15 (n = 5). (**E**) Representative EEG recordings presented as heat maps of frequency and amplitude data showing reduced seizure severity in mice treated with the CREB1 inhibitor 666-15. (**F**) Bar chart showing decreased EEG total power during SE in mice treated with the CREB1 inhibitor 666-15 (n = 9) when compared to vehicle-treated mice (n = 7). (**G**) Reduced transcript levels of the neuronal activity-regulated gene *c-Fos* in mice treated with the CREB1 inhibitor 666-15 when compared to vehicle-treated mice 8 h post-lorazepam injection (n = 5 per group). (**C-G**) Two-sided unpaired t-test. Data are mean ± S.E.M. * *P* < 0.05.

To test whether CREB1 has a role during seizure-induced pathology in the IAKA mouse model, mice were treated i.c.v. with the CREB1-specific inhibitor 666-15^30^ 10 min before IAKA injection and 1 h after treatment with lorazepam (**Fig. 4B**). Demonstrating CREB1 regulating the expression of genes linked to *Calcium signalling*, both *CamK4* and *Grm5* transcripts, down-regulated post-SE, were increased in mice treated with the CREB1 inhibitor 666-15 when compared to vehicle-injected mice 8 h post-SE (**Fig. 4C**). Suggesting a functional role for CREB1 during IAKA-induced SE, mice treated with the CREB1 inhibitor 666-15 experienced less severe seizures during SE as evidenced by a reduction in behavioural seizures (**Fig. 4D**), lower seizure total power (**Fig. 4E-F**) and lower transcript level of the neuronal activity-regulated gene *c-Fos* post-SE (**Fig. 4G**).

Thus, our results demonstrate that the IAKA model is a valid model to screen for and test novel anticonvulsive and antiepileptic drugs.

## Discussion

Here, by using a genome-wide gene expression analysis of the hippocampal transcriptome post-IAKA-induced SE, we demonstrate that the IAKA mouse model mimics closely molecular changes in the brain of patients with TLE once epilepsy is established and therefore represents a valid model for the testing of novel antiepileptogenic drug targets.

If we are to identify potential anti-epileptogenic and disease-modifying treatments for drug-resistant epilepsy, then the models we use must phenocopy the human pathophysiology. Large-scale changes in gene expression caused by injuries to the brain (*e.g.*, traumatic brain injury, SE) are widely recognized to contribute to the development of epilepsy.^5, 7, 40^ Using an arsenal of different experimental models of acquired epilepsy, we have now detailed knowledge of the disease-specific expression changes occurring in most of these models and of the molecular machinery driving these changes including transcriptional and post-transcriptional mechanisms.^6, 10, 41-43^ We have now extended this data by characterizing expression changes in the hippocampus of the IAKA mouse model^19^ during both SE and epilepsy.

A first result of our microarray analysis is that gene expression changes are much more prominent post-SE when compared to established epilepsy. This is in line with previous studies using models of SE induced via other chemoconvulsants.^8, 10^ Another result in agreement with previous studies is the fact that functional profiles are broadly unique to each condition.^8, 10^ While these findings are not surprising, they confirm the importance of transcriptional changes during the initial insult which may initiate a cascade of pathological changes leading eventually to the development of epilepsy. The reason for this initial surge in gene expression changes is most likely due to the increased neuronal excitability experienced during SE. Why SE and epilepsy lead to the dysregulation of a different gene pool remains elusive. The most obvious explanations include differences in seizure severity (SE *vs*. spontaneous seizure), time-point of tissue analysis relative to seizures and underlying pathology (*i.e.*, acute neurodegeneration *vs*. chronically diseased brain). Interestingly, genes dysregulated during both conditions are either up- or down-regulated, suggesting some common regulatory mechanisms during both SE and epilepsy.

Our pathway analysis of genes dysregulated in the IAKA model showed in particular genes linked to signalling pathways to be enriched among up-regulated genes and genes associated with metabolism and synaptic transmission among down-regulated genes post-SE. This is broadly in line with pathway analysis in other models (*e.g.*, MAPK family)^6, 10^ and suggests different stimuli activating similar pathways. During epilepsy, pathways most affected in the IKAK model included pathways involved in waste removal (up-regulated) and synaptic transmission (down-regulated). In particular, genes linked to lysosome activity were among the most affected genes. This is in good agreement with previous studies showing genes linked to the mTOR pathway to be commonly dysregulated during epilepsy.^8, 10^ Of note, autophagic/lysosomal system-related proteins are increasingly recognized to play an important role during epilepsy with drugs targeting autophagy reported to modulate seizures in several models.^44^

While we have amassed much data on gene expression in different models, there has been a paucity of cross-model comparisons. Here, we compared our findings in the IAKA mouse model with two commonly used mouse models to induce SE and epilepsy (*i.e.*, IHKA and i.p. pilocarpine). The main findings here were that gene expression changes are much more similar between both KA mouse models when compared to the pilocarpine model. An unexpected result was the almost complete overlap between both KA models, post-SE and during epilepsy. This is even more remarkable, as both models display different pathologies during and post-SE. This includes severe seizures during IAKA-induced SE with cell death mainly restricted to the ipsilateral CA3 subfield *vs*. mostly non-convulsive SE and widespread hippocampal neurodegeneration in the IHKA mouse model.^23^ In contrast, when we compared the IAKA model with the i.p. pilocarpine mouse model, similarities were less obvious, in particular post-SE. Thus, gene expression changes seem to be more dependent on chemoconvulsant used than pathology. It is, however, important to keep in mind that pilocarpine was delivered i.p. in contrast to the intracerebral delivery of KA in the two other models.

One of the main results of our study is the close correlation of gene expression in the KA models when compared to human TLE. This is an important finding as this strongly reinforces the rational for using these models to identify and test novel anti-epileptogenic drugs. Another important finding is that these similarities are restricted to established epilepsy and partly lost when we compared human tissue with mouse tissue collected shortly following SE, suggesting established epilepsy in mice is a much better model for TLE. Why the pilocarpine model shows less similarities with human TLE we do not know. The pilocarpine model is, however, associated with peripheral immune responses prior to the induction of SE and most likely reflects a mixture of an ischemic and excitotoxic insult.^45^ In line with a strong overlap in gene expression between the IAKA mouse model and TLE patients, pathway analysis of common dysregulated genes revealed similar pathways to be affected when compared to affected pathways during established epilepsy in the IAKA model. Most notably, calcium signalling is one of the most consistently affected pathways across models and patients which is consistent with previous reports.^6, 46, 47^ The effects of suppressed calcium signaling remain to be established. It is, however, tempting to speculate this being a protective mechanism reducing synaptic transmission and thereby hyperexcitability in brain tissue. Interestingly, calcium signalling was also suppressed in mice undergoing epileptic preconditioning.^24^

Finally, our IPA analysis predicted the cAMP responsive element binding protein CREB as one of the main up-stream transcription factors regulating gene transcription during SE and epilepsy. This is in line with previous findings identifying CREB as one of the main transcription factors in the pilocarpine mouse model.^10^ A role for CREB during seizures has been previously demonstrated with decreased levels of CREB leading to seizure reduction following pilocarpine in mice.^48^ Whether CREB contributes to seizures via the regulation of calcium signalling warrants, howver, further investigation. Interestingly, P53 was predicted as one of the main transcription factors driving gene transcription post-SE. P53 has previously been linked to seizure generation and cell death in the IAKA model.^49^ While our IPA analysis has focused primarily on transcription factors, other mechamiosns may also impact on mRNA levels such as microRNAs or mRNA polyadenylation among many others.^31, 41^

Possible limitations of our study include that gene expression has only been analyzed shortly post-SE and once epilepsy was already established omitting the seizure-free latent period. The latent period in the IAKA mouse model is, however, almost absent^23^ and the process of epileptogenesis is ongoing beyond the occurrence of the first epileptic seizure.^5^ Moreover, expression profiles have been analyzed using different analysis platforms (*e.g.*, microarray *vs*. sequencing) and cut-offs for analysis might be different. In addition, time-points analyzed relative to initial insult differ between animal models. We have, however, taken into account the different disease timelines of each model (*e.g.*, much longer latent period in the pilocarpine model compared to the IAKA model).^16, 45^ Moreover, some changes may have been masked by analyzing the whole hippocampus rather than subfields. Previous studies analysing gene expression in TLE patients have, however, used resected hippocampal tissue regardless of subfields.

In summary, our results demonstrate the IAKA mouse model closely mimicking human TLE thus providing strong rational for using the IAKA model to identify potential targets for disease-modifying treatments.

## Acknowledgments

This work was supported by funding from the Health Research Board HRA-POR-2015-1243 to T.E.; Science Foundation Ireland (17/CDA/4708 to T.E and co-funded under the European Regional Development Fund and by FutureNeuro industry partners 16/RC/3948 to D.C.H); from the H2020 Marie Skłodowska-Curie Actions Individual Fellowship (796600 to L.D-G and 753527 to E.B), from the European Union’s Horizon 2020 research and innovation programme under the Marie Sklodowska-Curie grant agreement (No. 766124 to T.E.). Further support was obtained from: PI2015-2/06-3&PI2018/06-1(ISCIII-CiberNed), SAF2015-65371-R (MINECO/AEI/FEDER, UE) and RTI2018-096322-B-I00 (MCIU/AEI/FEDER, UE) to J.J.L. A.P. was beneficiary of a Spark grant (SNSF) CRSK-3_190764/1. We thank the following core facilities: IRB-Functional Genomic and IRB-Bioinformatics/Biostatistics.

## Ethical Publication Statement

We confirm that we have read the Journal’s position on issues involved in ethical publication and affirm that this report is consistent with those guidelines.

## Figure Legends

**Supplementary Figure 1. Validation of differently expressed genes detected via microarray analysis.** (**A**) Log_2_ fold change of mRNA levels obtained by microarray analysis and (**B**) mRNA levels analysed by qRT-PCR and normalized to *β-Actin* of genes which are up-regulated post-SE and during epilepsy. Two-sided unpaired t-test. Data are mean ± S.E.M. * *P* < 0.05, ** *P* < 0.01, *** *P* < 0.001.

**Supplementary Figure 2. Additional comparisons of dysregulated genes between animal models of epilepsy and patients.** Graphs showing the comparison of dysregulated genes in TLE patients (hippocampus (left) and cortex (right)) with (**A**) the IHKA mouse model during epilepsy (15 days post-KA injection), (**B**) the IAKA mouse model following post-SE (8 h) and (**C**) the i.p. pilocarpine epilepsy model (6 weeks post-pilocarpine injection). Up-regulated genes common between two conditions are presented in yellow and down-regulated genes common between conditions in blue. Overlap: Representation Factor (RF) > 2 and *P* < 0.05, and dissimilar RF < 0.5 and P < 0.05. Hypergeometric test.

