## Supplementary figures and images for "High concordance between hippocampal transcriptome of the intraamygdala kainic acid model and human temporal lobe epilepsy"

### Supplementary Figures

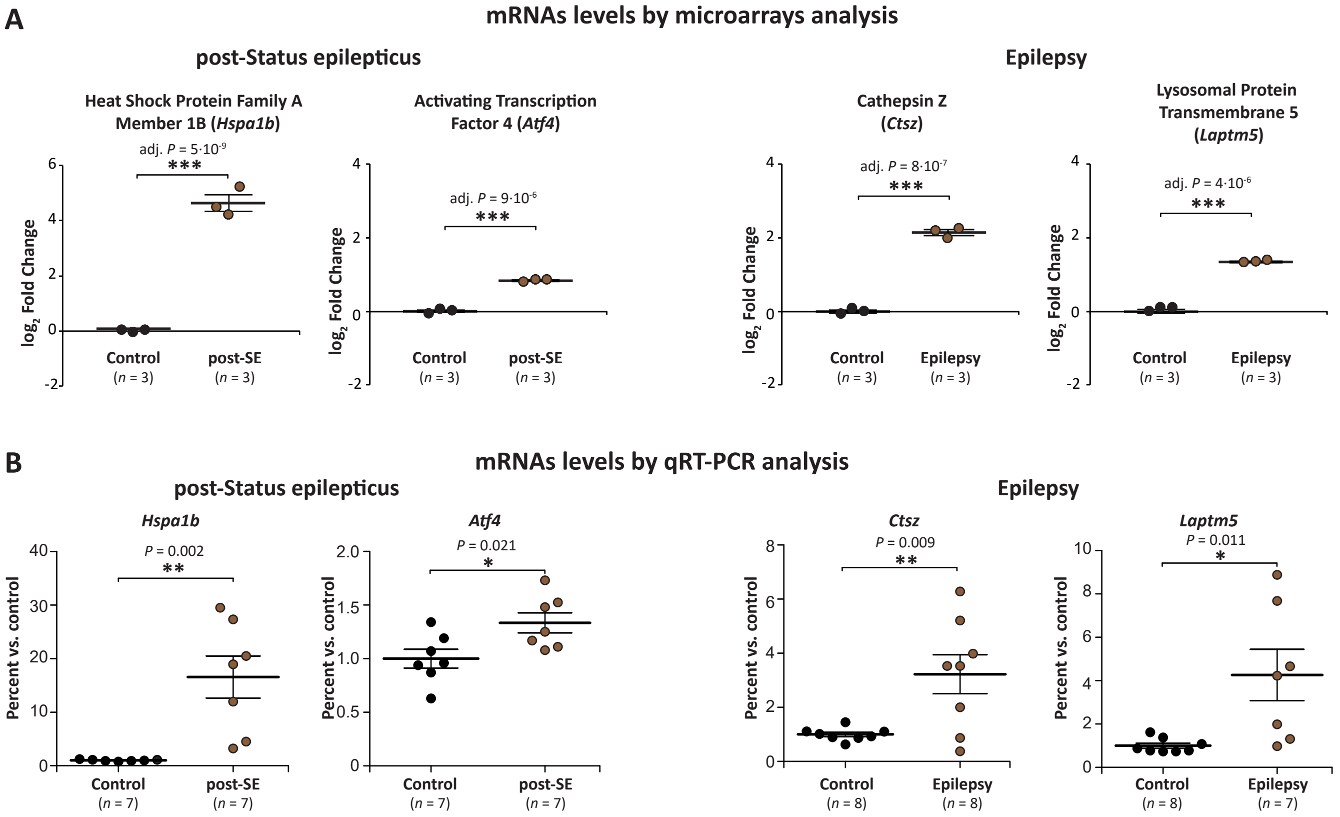


**Supplementary Figure 1**

#### Comparison of gene expression changes

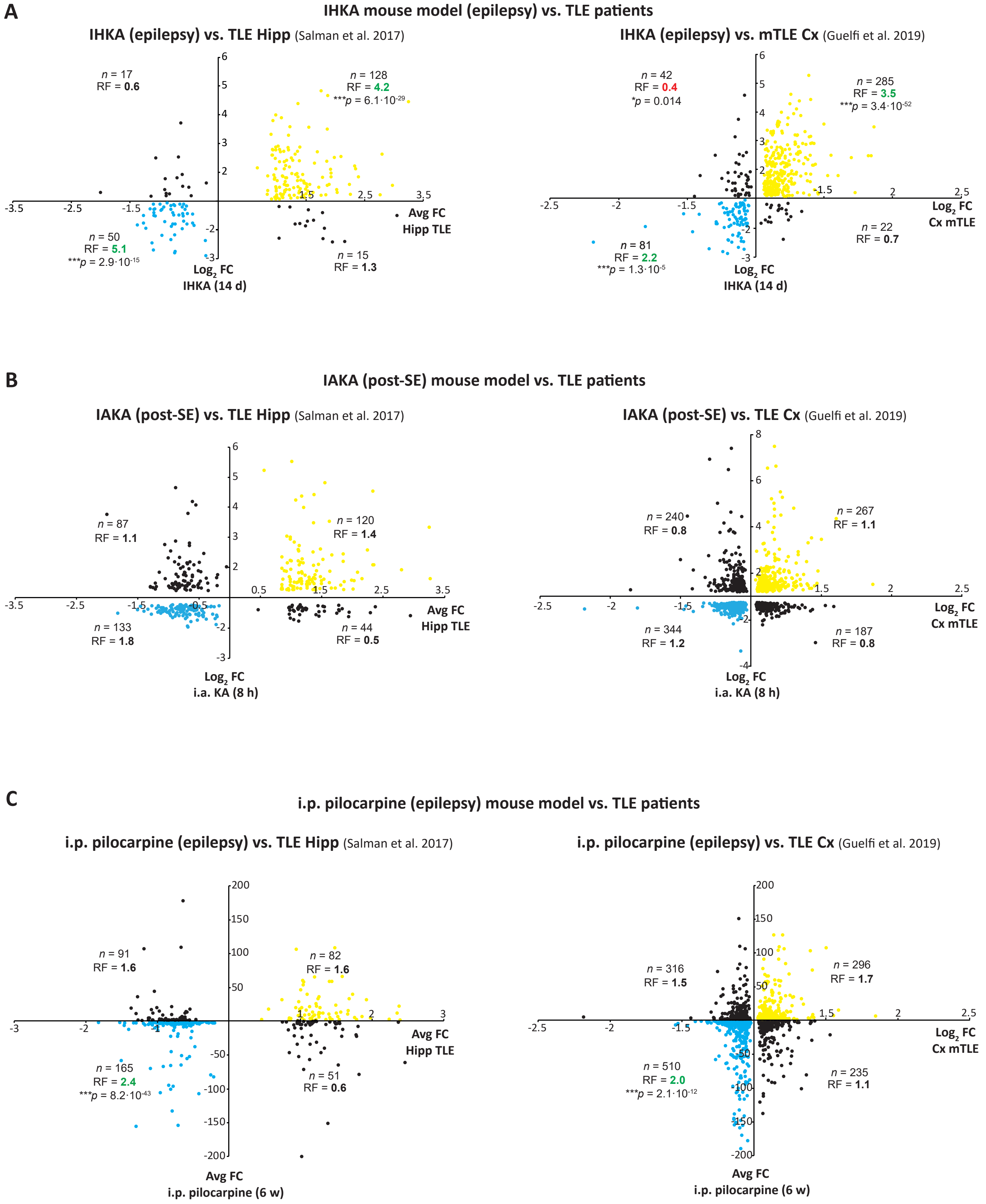


**Supplementary Figure 2**
